# Antigens in water-in-oil emulsion: a simple antigen extraction method for analysis and proof of equal antigen distribution in vaccination syringe after mixture

**DOI:** 10.1101/2020.01.22.916189

**Authors:** Michael Ghosh, Marion Gauger, Monika Denk, Hans-Georg Rammensee, Stefan Stevanović

**Author notes:** Author contributions: M.G., M.Ga., M.D., H.-G.R., S.S. designed research; M.G., M. Ga., M.D. performed research; M.G., M.Ga., and M.D. analyzed data; and M.G. and M. Ga. wrote the paper.

## Abstract

Therapeutic vaccination of antigens in oil emulsions is an approach becoming increasingly popular. Water-in-oil emulsions enable the formation of a depot, a slow passive antigen release and a decelerated antigen degradation. Furthermore, particularly advantageous for peptide vaccinations with a low clinical response rate is the increased immunogenicity and activation of CD8^+^ and CD4^+^ T-cells. Therefore, the use of personalized peptide vaccination cocktails in oil emulsions are tested in the treatment of cancer patients. An equal peptide distribution in the emulsion is striking for a successful depot effect to ensure the optimal efficacy and additionally, to enable reliable drug release analytics. First, the stability of the generated peptide-Montanide ISA™51 emulsion was demonstrated with a cocktail of heterogeneous peptides for 24 h in room temperature. Furthermore, we developed a simple peptide extraction method to investigate the equal peptide distribution in a Montanide ISA™51 emulsion in the syringe after mixing. Peptides were successfully extracted from a peptide-Montanide ISA™51 vaccination cocktail in sufficient volume for analysis and an equal peptide distribution in the syringe was verified via HPLC.

## Introduction

Immune checkpoint modulation has demonstrated the enormous potential of immunotherapies. Immune cells, especially T cells, which identify and eliminate cancer cells and lead to a long-lasting immunity are boosted. The T cell specificity is based on an interaction of the T cell receptor with Human Leukocyte Antigens (HLA) presented tumor specific peptides on the cancer cell membrane^1^. As a result, the use of multiple peptide antigens as vaccines is becoming an increasingly popular approach for cancer immunotherapy. Synthetic peptides are cheap, easy to produce in GMP quality, are storable in warehouses and therefore suitable for tailored vaccination cocktails ^2,3^. Peptide vaccination cocktails are used for individual and personalized treatment of cancer patients in vaccination studies and have demonstrated feasibility, safety and cancer regression, however also a low overall clinical response rate below 5 % ^3–5^.

Peptides are more or less not antigenic themselves and are dependent on the addition of adjuvants to enhance the efficacy and potency of the humoral and cellular anti-cancer responses ^3^. Besides the activation of antigen-presenting cells (APCs) and the induction of a long-term memory, some adjuvants protect the antigen from initial degradation, enable an even and long-lasting antigen release and support the antigen uptake ^3,6,7^. One of the most common adjuvants used for oil-in-water emulsions is Montanide ISA™51 (generally known: incomplete Freund’s adjuvant analogue). Clinical-grade Montanide ISA™51 has been widely used clinically in experimental peptide and protein-based cancer vaccines and as far as known it is safe and tolerable ^8^ except of reported sterile abscesses at the injection site ^9,10^. To this day, more than 40 vaccinations of synthetic peptides from the Wirkstoffpeptidlabor in Montanide ISA™51 emulsions did not lead to any severe system toxicities ^3,11–14^.

Since Montanide ISA™51 is so popular, it is all the more important to verify the homogeneity of the emulsion, so that an even antigen distribution is ensured in the syringe to achieve a successful depot effect and fully exploit the advantages of the combination with Montanide ISA™51. In order to analyze the peptide content with the HPLC, it is important to extract or separate the peptides of the water-in-oil emulsion. Hence, we developed an easy and fast extraction method, free from chemical contaminants, to enable the examination of the peptide distribution in the syringe.

## Methods

### Peptide synthesis

The peptide cocktails were synthesized based on the 9-fluorenylmethyloxycarbonyl/tert-butyl (Fmoc/tBu) strategy in GMP conditions and formulated into single-dose vials and released according to GMP at the Wirkstoffpeptidlabor of the Department of Immunology, University of Tübingen. *Peptide cocktails were formulated for NOA-16 (NCT02454634), IVAC-ALL-1 (NCT03559413) and GAPVAC-101 (NCT02149225)*^15^ *trials.*

### Peptide-Montanide ISA™51 emulsion

Mixing of a ten-peptide cocktail (Table 1) with adjuvant Montanide ISA™51 (SEPPIC) (50:50) generates a milky-white water-in-oil emulsion. First, the peptide cocktail solved in 33% DMSO (Dimethyl sulfoxide, Merck) and Montanide ISA™51 is defrozen for ten minutes. In the first syringe 600 µl peptide cocktail plus a similar volume of air and in the second syringe 600 µl Montanide ISA™51 plus a similar volume of air are taken up. The syringes are connected without air screwing a mixing chamber (Combifix adapter, B. Braun) on the first syringe, pressing out the remaining air. Then pressing out the remaining air of the second syringe and screwing it also to the mixing chamber.

**Table 1:** GMP-quality peptides used in the peptide-Montanide ISA™51 emulsions. A heterogenous mix of peptides with different lengths, masses, GRAVYs (Grand average of hydropathicity)^16^ and RT (Retention time with standard method), which were previously formulated as vaccination cocktail, were selected to investigate the emulsion stability. Peptide mass and GRAVY were calculated with https://web.expasy.org/cgi-bin/protparam/protparam.

| Cocktail | Peptide | Individual peptide concentration | Length | Mass [g/mol] | GRAVY |
| --- | --- | --- | --- | --- | --- |
| **1-peptide-cocktail** | IDH1R132H | 0.6 mg/ml | 20 | 2282.59 | -0.375 |
| **6-peptide-cocktail** | ALL1 | 1.0 mg/ml | 10 | 1090.28 | 0.510 |
|  | ALL2 | 1.0 mg/ml | 9 | 1159.31 | -0.500 |
|  | ALL3 | 1.0 mg/ml | 8 | 1018.14 | -0.713 |
|  | ALL4 | 1.0 mg/ml | 15 | 1803.09 | -0.467 |
|  | ALL5 | 1.0 mg/ml | 15 | 1913.13 | -1.887 |
|  | ALL6 | 1.0 mg/ml | 15 | 1790.05 | -1.060 |
| **10-peptide-cocktail** | GAPVAC1 | 0.8 mg/ml | 9 | 957.45 | 0.020 |
|  | GAPVAC2 | 0.8 mg/ml | 9 | 1090.48 | -1.256 |
|  | GAPVAC3 | 0.8 mg/ml | 9 | 1087.55 | -0.330 |
|  | GAPVAC4 | 0.8 mg/ml | 9 | 1098.48 | -0.320 |
|  | GAPVAC5 | 0.8 mg/ml | 10 | 1113.56 | 0.500 |
|  | GAPVAC6 | 0.8 mg/ml | 9 | 1117.59 | 0.210 |
|  | GAPVAC7 | 0.8 mg/ml | 9 | 1086.55 | 0.230 |
|  | GAPVAC8 | 0.8 mg/ml | 9 | 1095.57 | 0.430 |
|  | GAPVAC9 | 0.8 mg/ml | 9 | 1146.65 | 1.189 |
|  | GAPVAC10 | 0.8 mg/ml | 10 | 1132.66 | 1.570 |

To obtain a well-mixed emulsion the whole volume is pressed 20 times slowly between the two syringes for pre-emulsification, first. Subsequently, for the end emulsification the whole volume is shifted back and forth 40 times quickly until a white stable emulsion is generated.

In order to verify the stability of the emulsion, the sample is pressed into one syringe and the Combifix adapter is replaced with a needle. One drop of the emulsion is placed in water and another drop on a microscope slide with 10% inclination. If the drop separates in two phases in water or is running down the slide, the emulsion is not stable and the mixing steps need to be repeated.

### Extraction of peptides in water-in-oil emulsion

To analyze the equal distribution of the emulsion in the syringe the emulsion is aliquoted in three equal parts (front, middle, back). Samples are frozen for at least 10 min at −80°C and then immediately centrifuged for 90 min at 4°C and 13000 rpm (Heraeus™ Biofuge fresco, Thermo Scientific). After centrifugation three different stable phases are obtained. The upper phase contains majorly Montanide ISA™51 (n=1.4625), the center phase a Montanide ISA™51 emulsion with DMSO (solid phase without refraction index), only obtained after successful mixing, and the lowest phase peptides solved in water (n=1.3350), assumed based on the refraction indexes. From the lower phase 40 µl clear solution were aliquoted and frozen at −80°C until analysis.

### HPLC and LC-MS/MS analysis

Analytical HPLC peptide analysis was performed on an e2695 Alliance System (Waters). 10 µl clear lower phase analyte were injected for reverse phase separation on a 3 x 250 mm separation column (Multospher 120 RP 18-5 µ) and UV-light absorption was measured at 220 nm. Peptide separation was performed at 30 °C and a flow rate of 0,2 ml/min applying a gradient ranging from 5-65 % of acetonitrile.

The peptide sequences of peptide peaks identified in the HPLC chromatogram were verified in each experiment using a LC-MS/MS (data not shown). LC-MS/MS peptide analysis was performed on a linear trap quadrupole (LTQ) Orbitrap XL mass spectrometer (Thermo Fisher) and coupled to an Ultimate 3000 RSLC Nano UHPLC System (Dionex). Peptide separation was performed at 30 °C and a flow rate of 5 µl/min on a 300 µm 15 cm separation column (Acclaim PepMap RSLC; Thermo Fisher) applying a gradient ranging from 4 to 44.0% of acetonitrile. Samples were analyzed on an LTQ Orbitrap XL using a top three CID (collision-induced dissociation) method.

## Results

### Peptide extraction protocol and experimental setting

In order to investigate the peptide distribution in the final peptide-Montanide ISA™51 emulsion intended for vaccination, we performed reconstitutions of GMP synthesized peptide cocktails and clinical grade Montanide ISA™51 and extracted the peptides according to protocol in Figure 1. In order to obtain a homogenous mixture, a low pace pre-emulsification and subsequent high-speed emulsification was performed as described in the method part. For HPLC and LC-MS/MS analysis, the reconstitution was aliquoted three times, frozen and centrifuged at maximum speed. From the resulting three stable phases 40 µl of the water-peptide solution of the lowest phase was extracted for analysis. There are several peptide extraction kits available, however as the homogenous peptide-Montanide ISA™51 mixture is only constituted with effort, we were able to successfully extract peptides based on simple freezing and centrifugation. Besides the simplicity another advantage of this extraction is the lack of by-products interfering with the HPLC and LC-MS/MS analysis.

**Figure 1:**
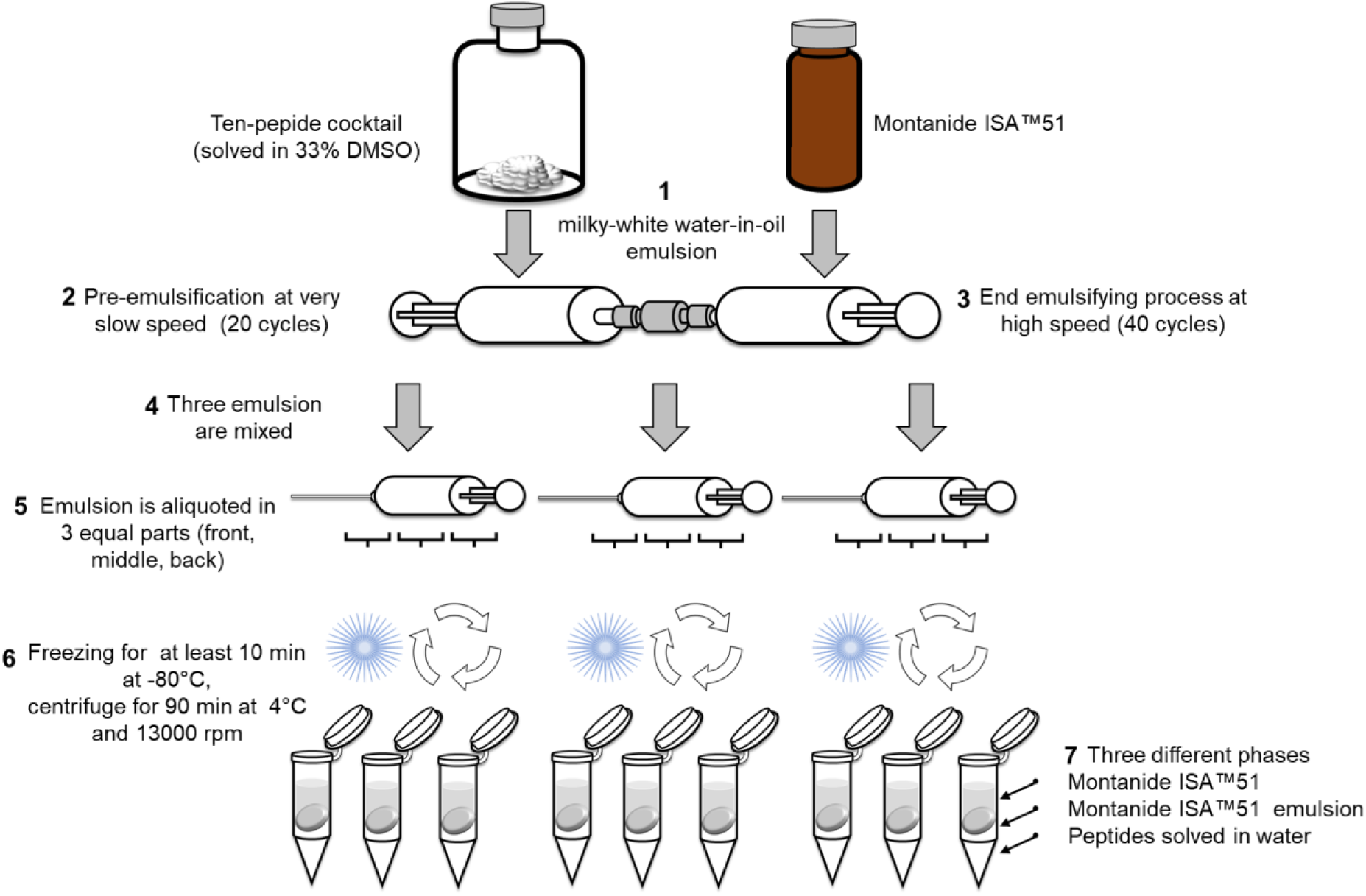
Overview of the emulsification and the peptide extraction protocol from the peptide-Montanide ISA™51 emulsion and aliquots selected for HPLC and LC-MS/MS analysis.

### Emulsion stability

To guarantee the quality of the emulsion according to the protocol for further experiments and later application in practice, the emulsion should remain stable for at least 24 hours at room temperature. Here, we performed the reconstitution according to the protocol for one, six and ten peptide cocktails, which were previously used in vaccination trials *(Table 1; NOA-21: NCT03893903, IVAC-ALL-1: NCT03559413, GAPVAC: NCT02149225).*

Three peptide-Montanide ISA™51 emulsions of each cocktail were prepared from three different persons and stored for 24 h at room temperature. Each product was observed and photographed after 0, 1, 2, 3, 4, 18 and 24 h after mixing. Immediately after the mixing and after 24 h one drop of the emulsion was placed through a needle in water and another drop on a microscope slide with 10% inclination. No drop separated in two phases and the emulsion was stable as white emulsion for 24 h (Figure 2).

**Figure 2:**
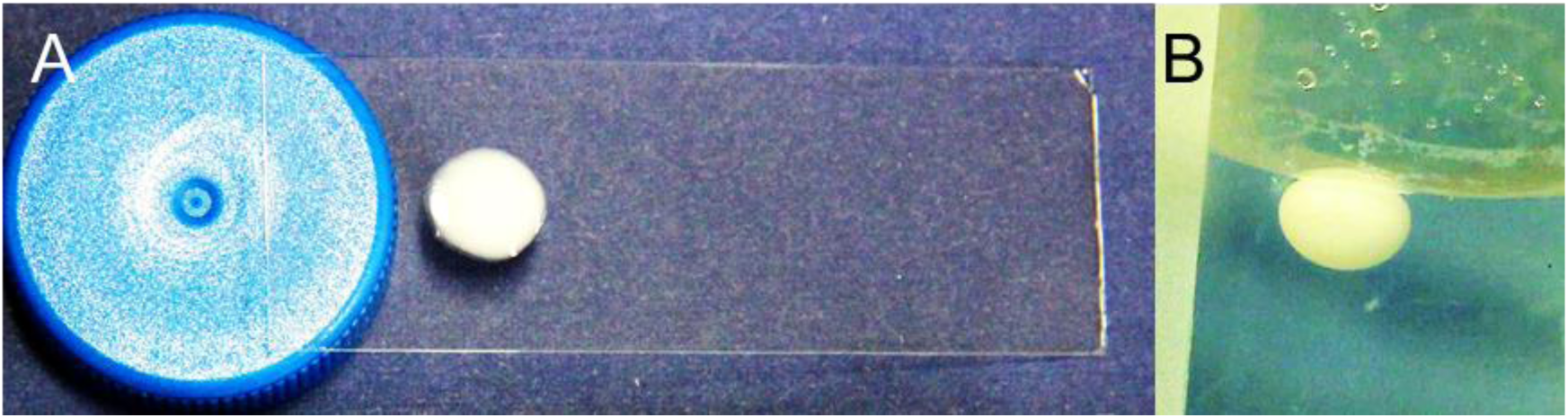
Quality control of the peptide-Montanide ISA™51 emulsions. **A**: One drop of the emulsion was placed after 0 and 24 h on a microscope slide with 10% inclination and should not separate in oil and water phases (**A**) and another drop was added in water and should remain compact and do not dissolve (**B**).

### Similar peptide distribution in the syringe

For the investigation of the peptide distribution in the syringe, three fresh peptide-Montanide ISA™51 emulsions were prepared according to protocol (Figure 1). The front, middle and rear thirds of the syringe were aliquoted into reaction tubes. The peptides dissolved in water in the lowest phase were analyzed with HPLC and LC-MS/MS. Chromatograms of the performed experiments are given in Supplemental Figure S1 (extracted peptide cocktail and initial peptide cocktail). In Figure 3A the area of each peptide in the three aliquots of the three emulsions is indicated. For all ten peptides of the cocktail the peptide area is similar for the three aliquots with almost no deviation. In a comparison of each peptide quantity in the three syringes, the relative standard deviation of each peptide area is below 3 %RSD, demonstrating the homogeneity of the emulsion in each part of the syringe (Figure 3B). The retention times stay similar between the aliquots as well as the areas did (Figure 4A). Also, between the peptide RTs of the whole syringes there is a low deviation with a maximal STD of 0.05 min (Figure 4B). Since the extraction protocol was supposed to be as simple, fast and reproducible as possible, the extracted peptide amount of the three peptides GAPVAC1, GAPVAC10 and GAPVAC2, which peaks are clearly separated in the chromatogram, were compared with the original peptide cocktail (Figure 5). The % area loss of the three peptides depends strongly on the individual peptide and reveals a higher deviation than the peak height of the three peptides. After extraction of the peptides GAPVAC10 and GAPVAC2 only 20% were lost of their original area, whereas in the case of GAPVAC1 40% less was extracted. Interestingly, the three peptides have different GRAVYs, GAPVAC10 (GRAVY=1.570) is highly hydrophobic and elutes last, whereas GAPVAC2 (GRAVY=-1.256) is highly hydrophilic and elutes second and GAPVAC1 (GRAVY=0.020) has a GRAVY in between and elutes first of all ten peptides (Table 1, Figure 4A). In comparison with the initial peptide cocktail all three peptides elute later with a difference of 0,08 to 0,42 minutes delay (Figure 5).

**Figure 3:**
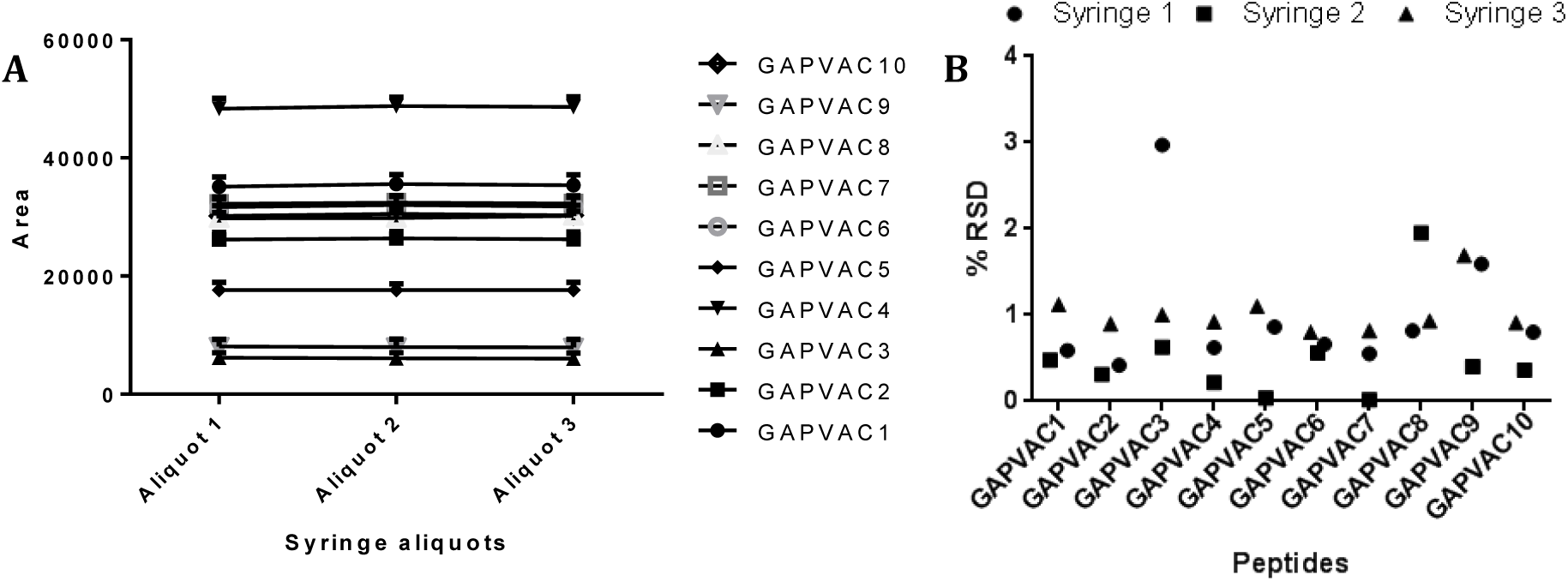
Changes in peptide quantity (area) after extraction from the peptide-Montanide ISA™51 emulsion. **A**: Mean peptide areas and deviation of the ten peptides in the peptide-extract of the three aliquots averaged for the three syringes. **B**: Percental RSD of the peptide area in the peptide-extract of the three aliquots of each syringe.

**Figure 4:**
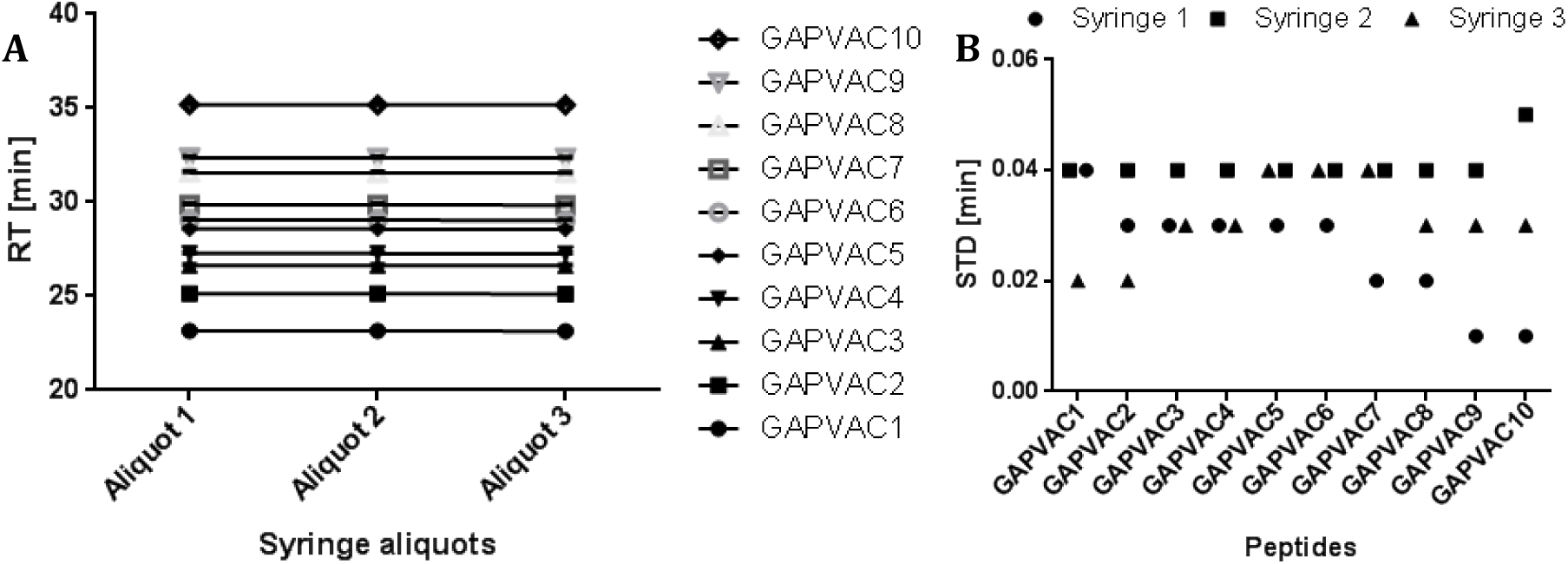
Changes in peptide RT after extraction from the peptide-Montanide ISA™51 emulsion. **A**: Mean retention times and deviation of the ten peptides in the peptide-extract of the three aliquots averaged for the three syringes. **B**: Deviation of the peptide retention times in the peptide-extract of the three aliquots of each syringe.

**Figure 5:**
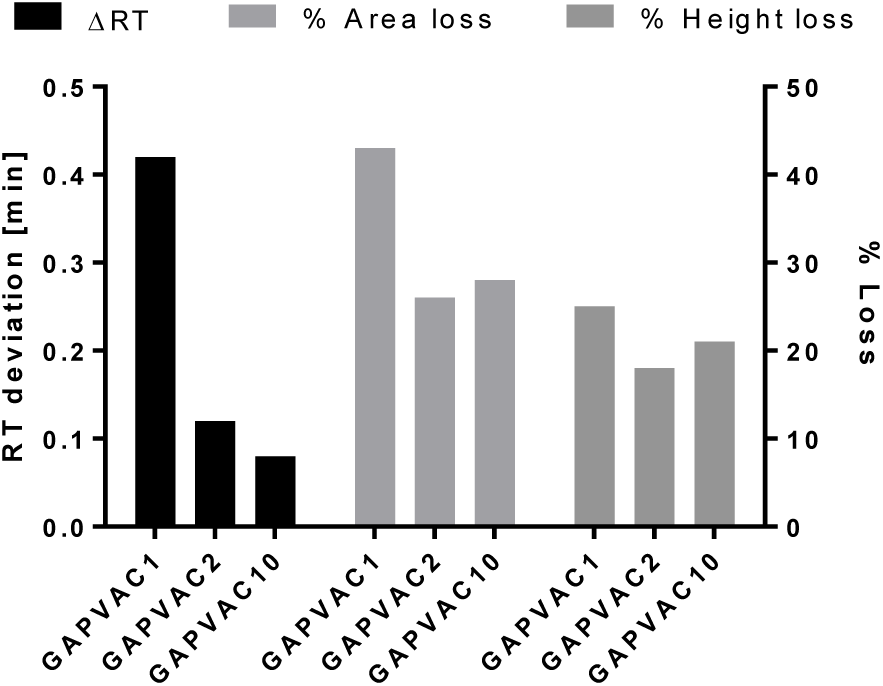
Changes of the peptide chromatogram RT, area and height after extraction from the peptide-Montanide ISA™51 emulsion. Comparison of a consecutive measurement of 10 µl of the initial peptide cocktail and 10 µl of the extracted lower phase after emulsion. The deviating retention time (left y axis), percental area loss and height loss (right y axis) in the chromatogram after extraction are indicated for the three peptides GAPVAC1, GAPVAC10 and GAPVAC2.

## Discussion

Montanide ISA™51 as adjuvant intended to be used for therapeutic vaccinations is known to be a potent adjuvant inducing CD4+ and CD8+ T cell responses. We developed a simple and by-product free protocol, which enabled the extraction of ten peptides in water-in-oil emulsions with Montanide ISA™51 by freezing and centrifugation. The emulsion was separated into three phases, which were assumed from the top as Montanide ISA™51, a Montanide ISA™51 emulsion with DMSO, and peptides solved in water. This simple protocol is probably limited for hydrophobic peptides, however in the ten-peptide cocktail used for this study no unambiguous correlation of the recovered peptide content and of the peptide GRAVY was recognizable. We could show, the distribution of peptides after the emulsification process is equal (in the whole syringe) which is striking for a homogeneous patient product, a successful depot effect with optimal efficacy and the analysis by HPLC and therefore for the quality control of vaccine peptide cocktails. When checking the emulsion after mixing by drop control, there will be no loss due to uneven distribution. After extraction a so far unpredictable individual peptide loss and a shift in the retention time needs to be considered. The retention time might be deviating due to remaining Montanide ISA™51 lipids in the extract, leading to a different peptide column matrix interaction. In this study we confirmed the homogenous mixture of the peptide-Montanide ISA™51 emulsions in the syringe, provided a simple protocol for a successful mixture, extraction and verification to support further clinical trials and the next generation cancer vaccines with Montanide ISA™51 combinations to achieve synergistic effects.

## Abbreviations

AcN: Acetonitrile
APC: Antigen-presenting cell
DMSO: Dimethyl sulfoxide
GMP: Good manufacturing practice
HPLC: High performance liquid chromatography
LC-MS/MS: Liquid chromatography tandem mass spectrometry
STD: Standard deviation

## Acknowledgements

This work was supported by the German Cancer Consortium (DKTK) and the Natural and Medical Sciences Institute at the University of Tübingen NMI. We thank the Wirkstoffpeptidlabor, especially Patricia Hrstic, Ulrich Wulle and Nicole Bauer for expert peptide synthesis and quality control.

## References

1. Freudenmann, L. K., Marcu, A. & Stevanović, S. Mapping the tumour human leukocyte antigen (HLA) ligandome by mass spectrometry. Immunology 154, 331–345 (2018).

2. Rammensee, H.-G., Weinschenk, T., Gouttefangeas, C. & Stevanović, S. Towards patient-specific tumor antigen selection for vaccination. Immunol. Rev. 188, 164–76 (2002).

3. Gouttefangeas, C. & Rammensee, H.-G. Personalized cancer vaccines: adjuvants are important, too. Cancer Immunol. Immunother. 67, 1911–1918 (2018).

4. Obeid, J., Hu, Y. & Slingluff, C. L. Vaccines, Adjuvants, and Dendritic Cell Activators—Current Status and Future Challenges. Semin. Oncol. 42, 549–561 (2015).

5. Rosenberg, S. A., Yang, J. C. & Restifo, N. P. Cancer immunotherapy: moving beyond current vaccines. Nat. Med. 10, 909–15 (2004).

6. Reed, S. G., Orr, M. T. & Fox, C. B. Key roles of adjuvants in modern vaccines. Nat. Med. 19, 1597–1608 (2013).

7. De Gregorio, E., Caproni, E. & Ulmer, J. B. Vaccine Adjuvants: Mode of Action. Front. Immunol. 4, 214 (2013).

8. Chiang, C. L.-L., Kandalaft, L. E. & Coukos, G. Adjuvants for Enhancing the Immunogenicity of Whole Tumor Cell Vaccines. Int. Rev. Immunol. 30, 150–182 (2011).

9. van Doorn, E., Liu, H., Huckriede, A. & Hak, E. Safety and tolerability evaluation of the use of Montanide ISA™51 as vaccine adjuvant: A systematic review. Hum. Vaccin. Immunother. 12, 159–169 (2016).

10. Graham, B. S., McElrath, M. J., Keefer, M. C., Rybczyk, K., Berger, D., Weinhold, K. J., Ottinger, J., Ferarri, G., Montefiori, D. C., Stablein, D., Smith, C., Ginsberg, R., Eldridge, J., Duerr, A., Fast, P., Haynes, B. F. & AIDS Vaccine Evaluation Group. Immunization with Cocktail of HIV-Derived Peptides in Montanide ISA-51 Is Immunogenic, but Causes Sterile Abscesses and Unacceptable Reactogenicity. PLoS One 5, e11995 (2010).

11. Löffler, M. W., Chandran, P. A., Laske, K., Schroeder, C., Bonzheim, I., Walzer, M., Hilke, F. J., Trautwein, N., Kowalewski, D. J., Schuster, H., Günder, M., Carcamo Yañez, V. A., Mohr, C., Sturm, M., Nguyen, H.-P., Riess, O., Bauer, P., Nahnsen, S., Nadalin, S., Zieker, D., Glatzle, J., Thiel, K., Schneiderhan-Marra, N., Clasen, S., Bösmüller, H., Fend, F., Kohlbacher, O., Gouttefangeas, C., Stevanović, S., Königsrainer, A. & Rammensee, H.-G. Personalized peptide vaccine-induced immune response associated with long-term survival of a metastatic cholangiocarcinoma patient. J. Hepatol. 65, 849–855 (2016).

12. Feyerabend, S., Stevanovic, S., Gouttefangeas, C., Wernet, D., Hennenlotter, J., Bedke, J., Dietz, K., Pascolo, S., Kuczyk, M., Rammensee, H.-G. & Stenzl, A. Novel multi-peptide vaccination in Hla-A2+ hormone sensitive patients with biochemical relapse of prostate cancer. Prostate 69, 917–27 (2009).

13. Rausch, S., Gouttefangeas, C., Hennenlotter, J., Laske, K., Walter, K., Feyerabend, S., Chandran, P. A., Kruck, S., Singh-Jasuja, H., Frick, A., Kröger, N., Stevanović, S., Stenzl, A., Rammensee, H.-G. & Bedke, J. Results of a Phase 1/2 Study in Metastatic Renal Cell Carcinoma Patients Treated with a Patient-specific Adjuvant Multi-peptide Vaccine after Resection of Metastases. Eur. Urol. Focus (2017). doi: 10.1016/j.euf.2017.09.009

14. Rammensee, H.-G., Wiesmüller, K.-H., Chandran, P. A., Zelba, H., Rusch, E., Gouttefangeas, C., Kowalewski, D. J., Di Marco, M., Haen, S. P., Walz, J. S., Gloria, Y. C., Bödder, J., Schertel, J.-M., Tunger, A., Müller, L., Kießler, M., Wehner, R., Schmitz, M., Jakobi, M., Schneiderhan-Marra, N., Klein, R., Laske, K., Artzner, K., Backert, L., Schuster, H., Schwenck, J., Weber, A. N. R., Pichler, B. J., Kneilling, M., la Fougère, C., Forchhammer, S., Metzler, G., Bauer, J., Weide, B., Schippert, W., Stevanović, S. & Löffler, M. W. A new synthetic toll-like receptor 1/2 ligand is an efficient adjuvant for peptide vaccination in a human volunteer. J. Immunother. Cancer 7, 307 (2019).

15. Hilf, N., Kuttruff-Coqui, S., Frenzel, K., Bukur, V., Stevanović, S., Gouttefangeas, C., Platten, M., Tabatabai, G., Dutoit, V., van der Burg, S. H., thor Straten, P., Martínez-Ricarte, F., Ponsati, B., Okada, H., Lassen, U., Admon, A., Ottensmeier, C. H., Ulges, A., Kreiter, S., von Deimling, A., Skardelly, M., Migliorini, D., Kroep, J. R., Idorn, M., Rodon, J., Piró, J., Poulsen, H. S., Shraibman, B., McCann, K., Mendrzyk, R., Löwer, M., Stieglbauer, M., Britten, C. M., Capper, D., Welters, M. J. P., Sahuquillo, J., Kiesel, K., Derhovanessian, E., Rusch, E., Bunse, L., Song, C., Heesch, S., Wagner, C., Kemmer-Brück, A., Ludwig, J., Castle, J. C., Schoor, O., Tadmor, A. D., Green, E., Fritsche, J., Meyer, M., Pawlowski, N., Dorner, S., Hoffgaard, F., Rössler, B., Maurer, D., Weinschenk, T., Reinhardt, C., Huber, C., Rammensee, H.-G., Singh-Jasuja, H., Sahin, U., Dietrich, P.-Y. & Wick, W. Actively personalized vaccination trial for newly diagnosed glioblastoma. Nature 565, 240–245 (2019).

16. Kyte, J. & Doolittle, R. F. A simple method for displaying the hydropathic character of a protein. J. Mol. Biol. 157, 105–132 (1982).

