## Supplemental Figure S1 for "Antigens in water-in-oil emulsion: a simple antigen extraction method for analysis and proof of equal antigen distribution in vaccination syringe after mixture"

### Initial ten-peptide cocktail

#### Ten-peptide cocktail 1411

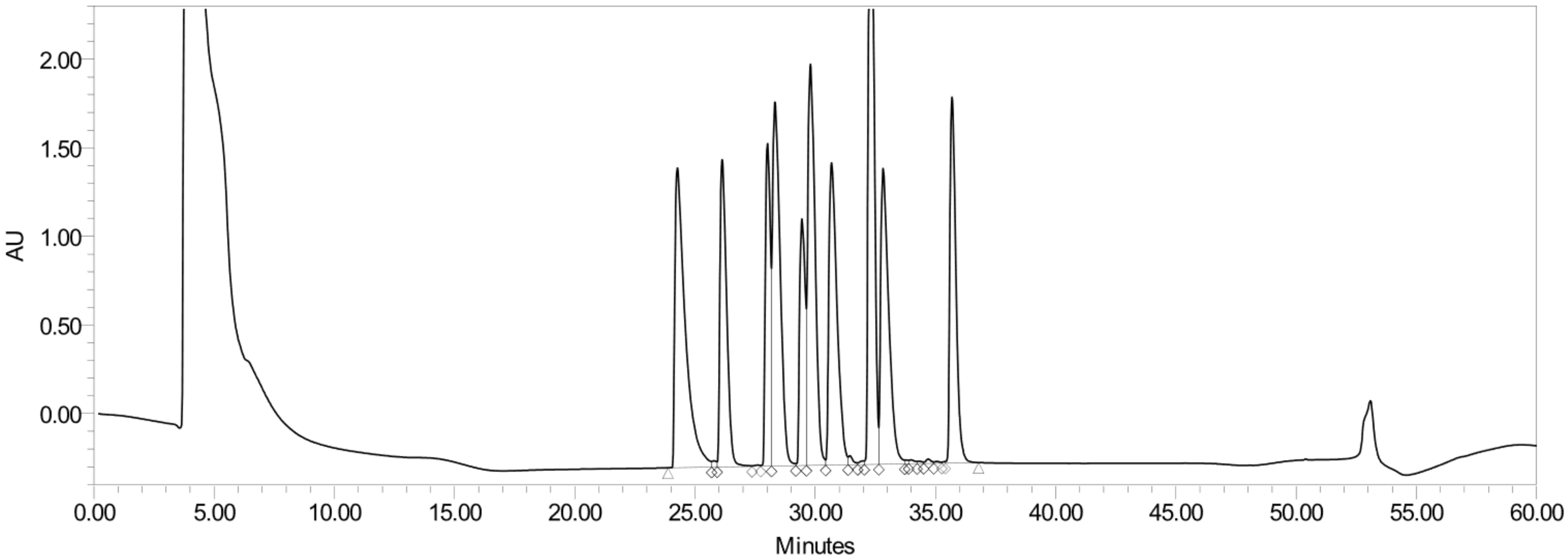

### Peptides extracted from the ten-peptide cocktail Montanide ISA™ 51 emulsion

#### Extracted lower phase 1.1

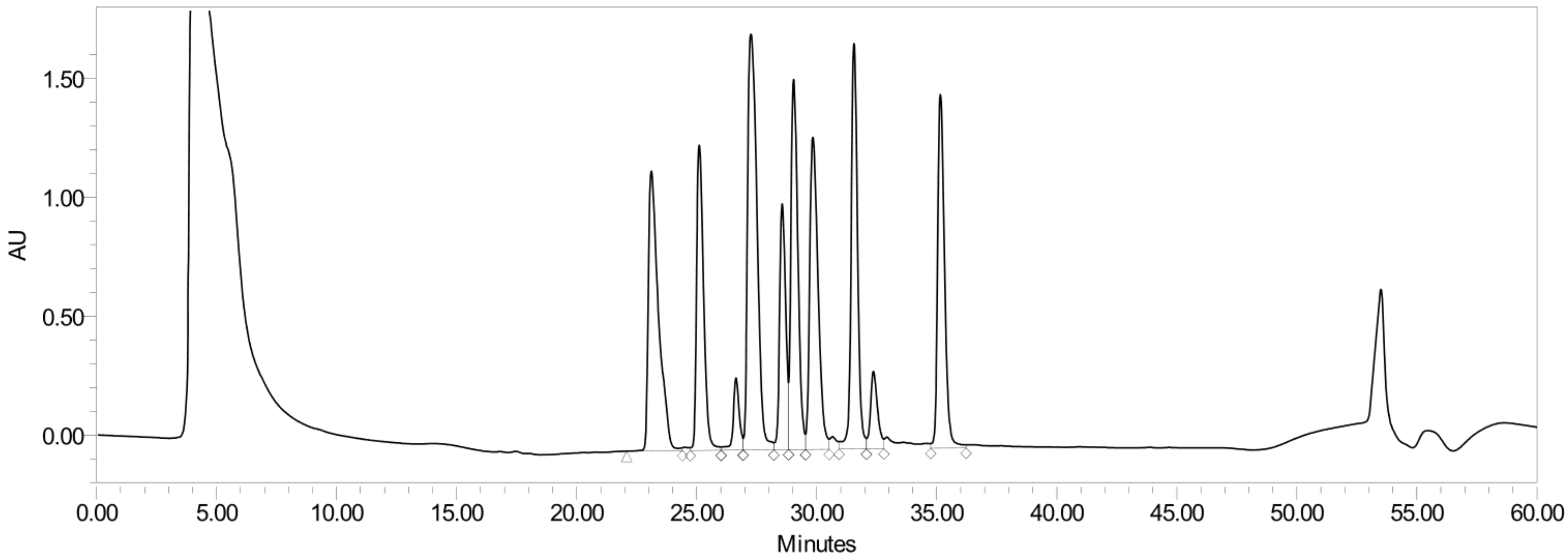

### Peptides extracted from the ten-peptide cocktail Montanide ISA$^{TM}$ 51 emulsion

#### Extracted lower phase 1.2

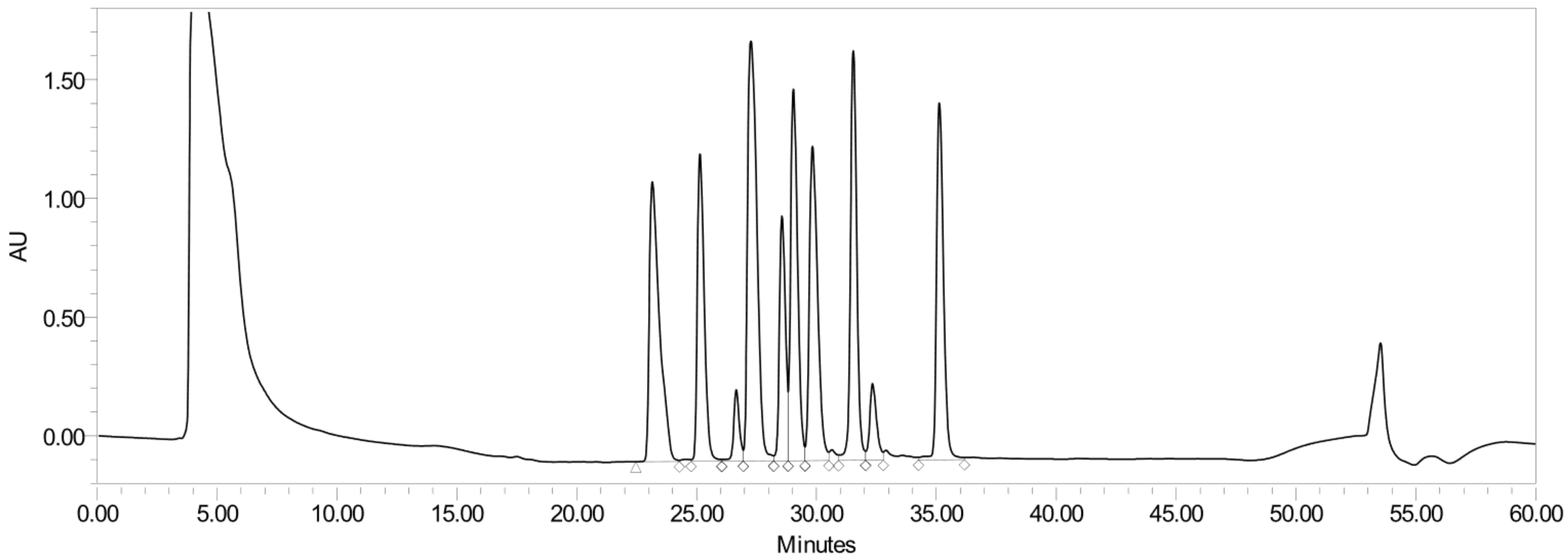

### Peptides extracted from the ten-peptide cocktail Montanide ISA$^{TM}$ 51 emulsion

#### Extracted lower phase 1.3

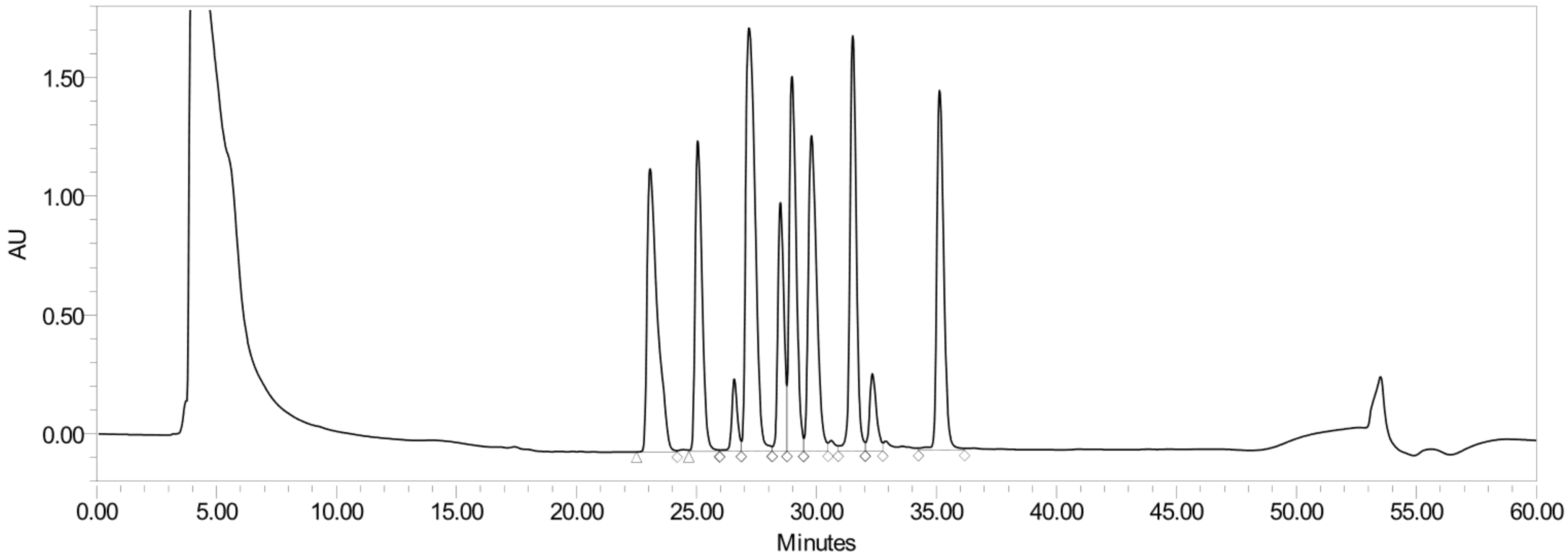

### Peptides extracted from the ten-peptide cocktail Montanide ISA[TM] 51 emulsion

#### Extracted lower phase 2.1

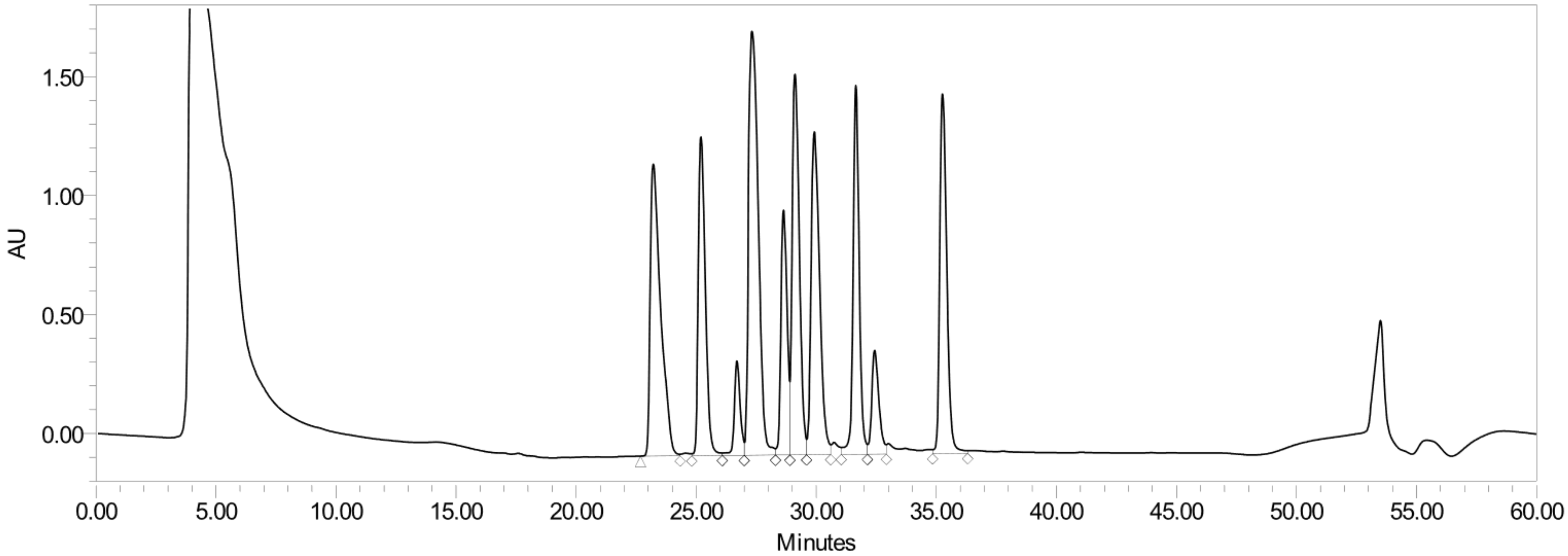

### Peptides extracted from the ten-peptide cocktail Montanide ISA$^{TM}$ 51 emulsion

#### Extracted lower phase 2.2

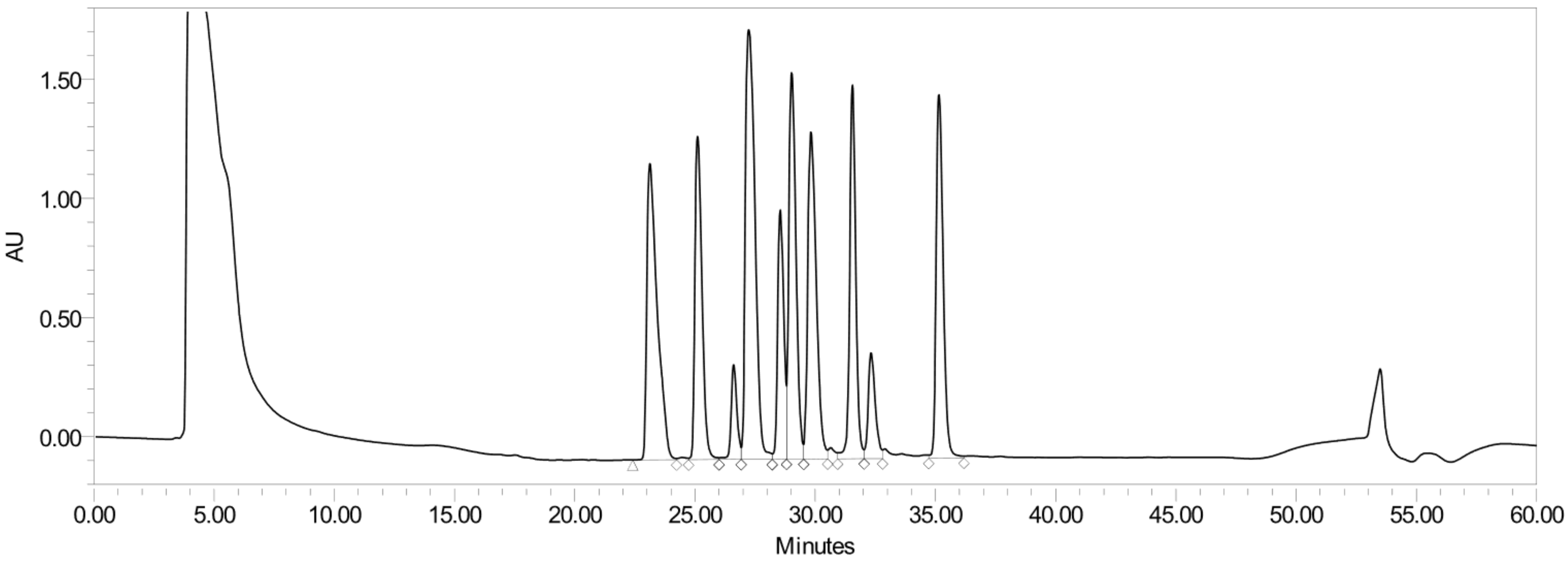

### Peptides extracted from the ten-peptide cocktail Montanide ISA$^{TM}$ 51 emulsion

#### Extracted lower phase 2.3

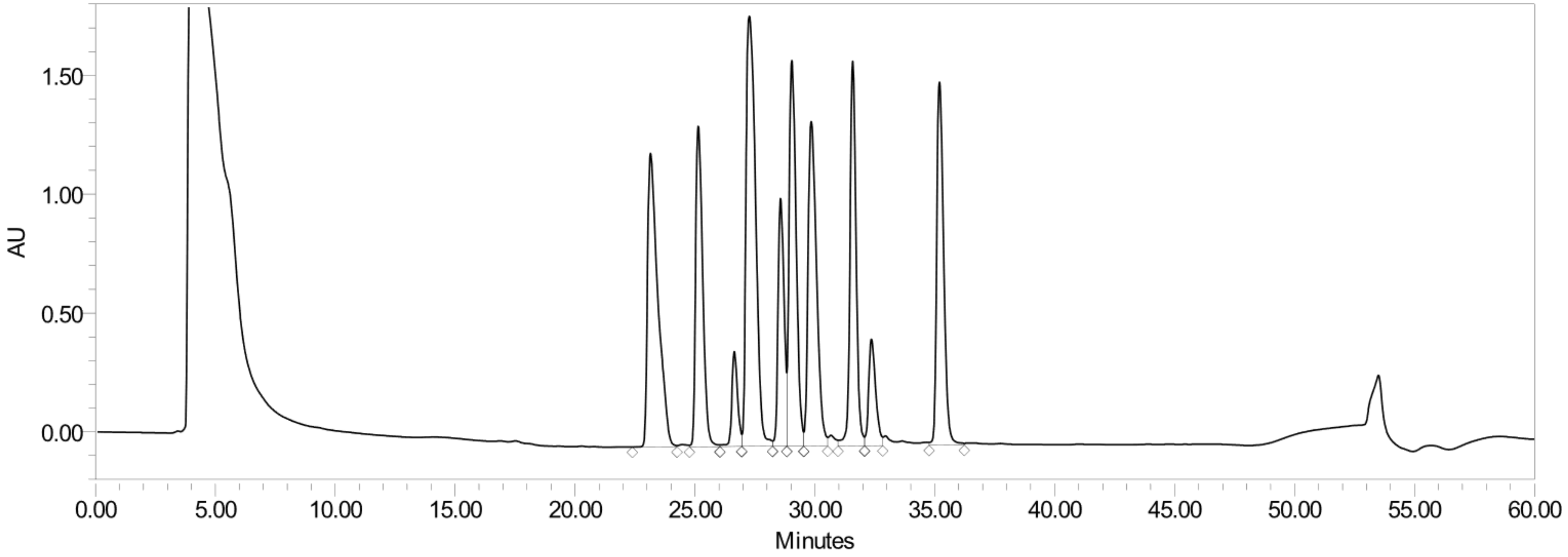

### Peptides extracted from the ten-peptide cocktail Montanide ISA$^{TM}$ 51 emulsion

#### Extracted lower phase 3.1

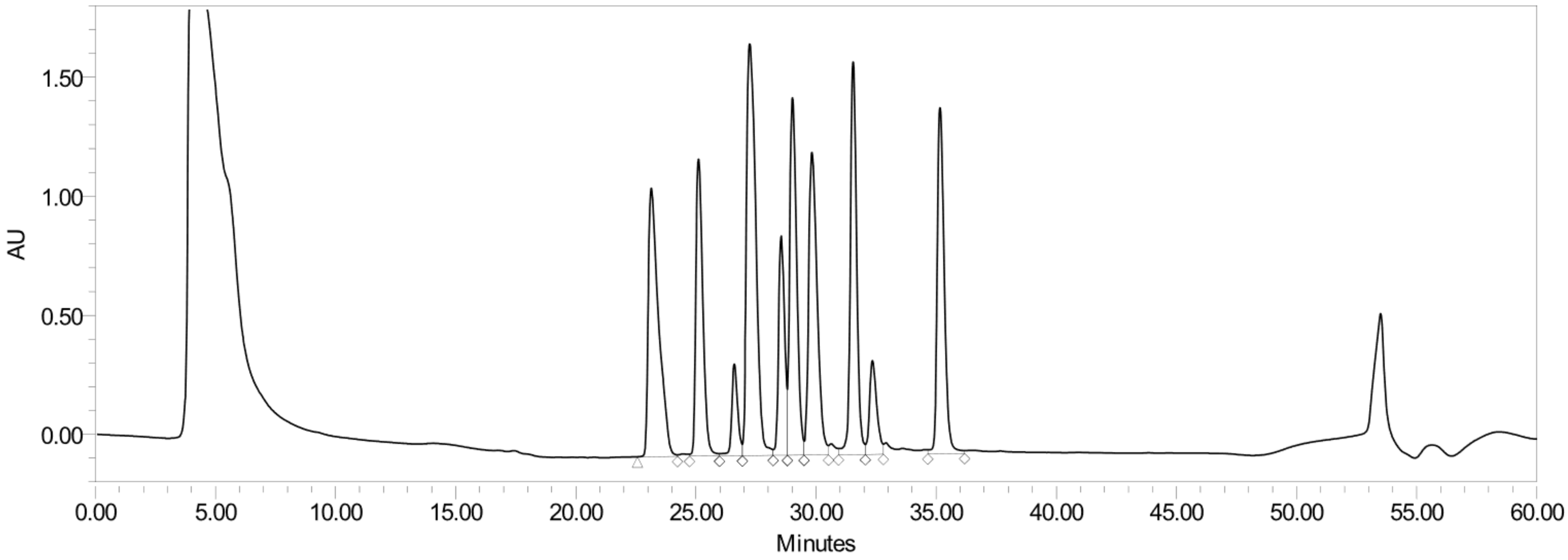

### Peptides extracted from the ten-peptide cocktail Montanide ISA$^{TM}$ 51 emulsion

#### Extracted lower phase 3.2

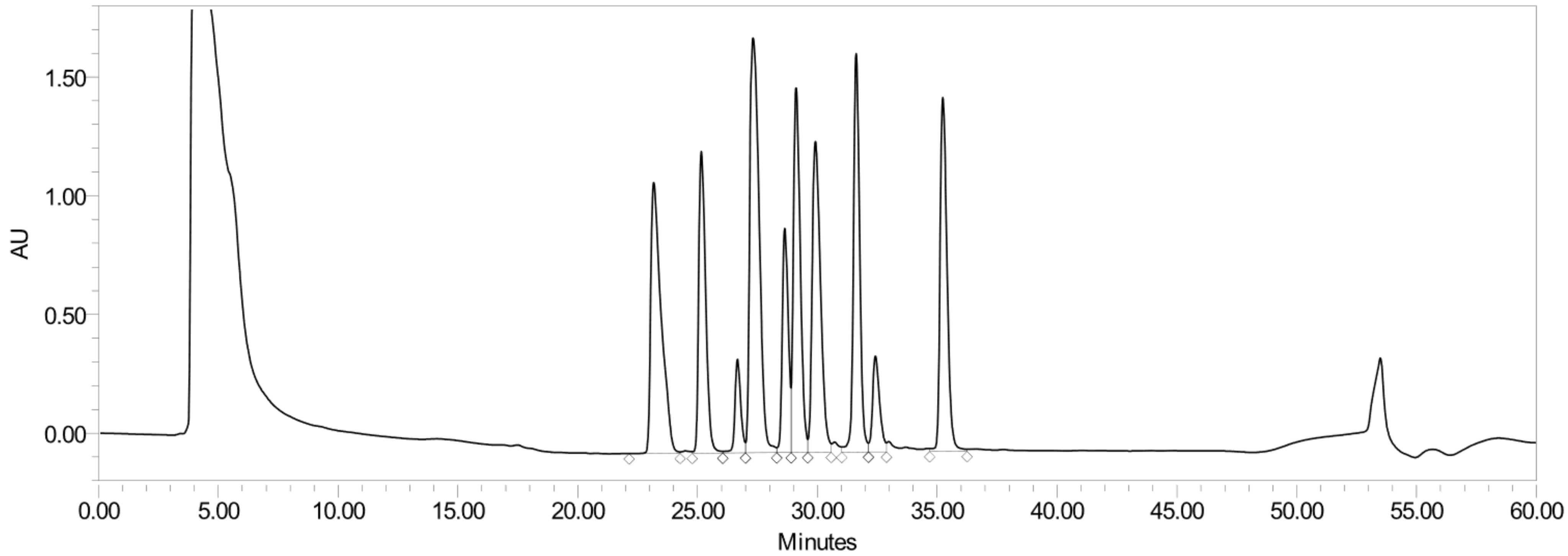

### Peptides extracted from the ten-peptide cocktail Montanide ISA$^{TM}$ 51 emulsion

#### Extracted lower phase 3.3

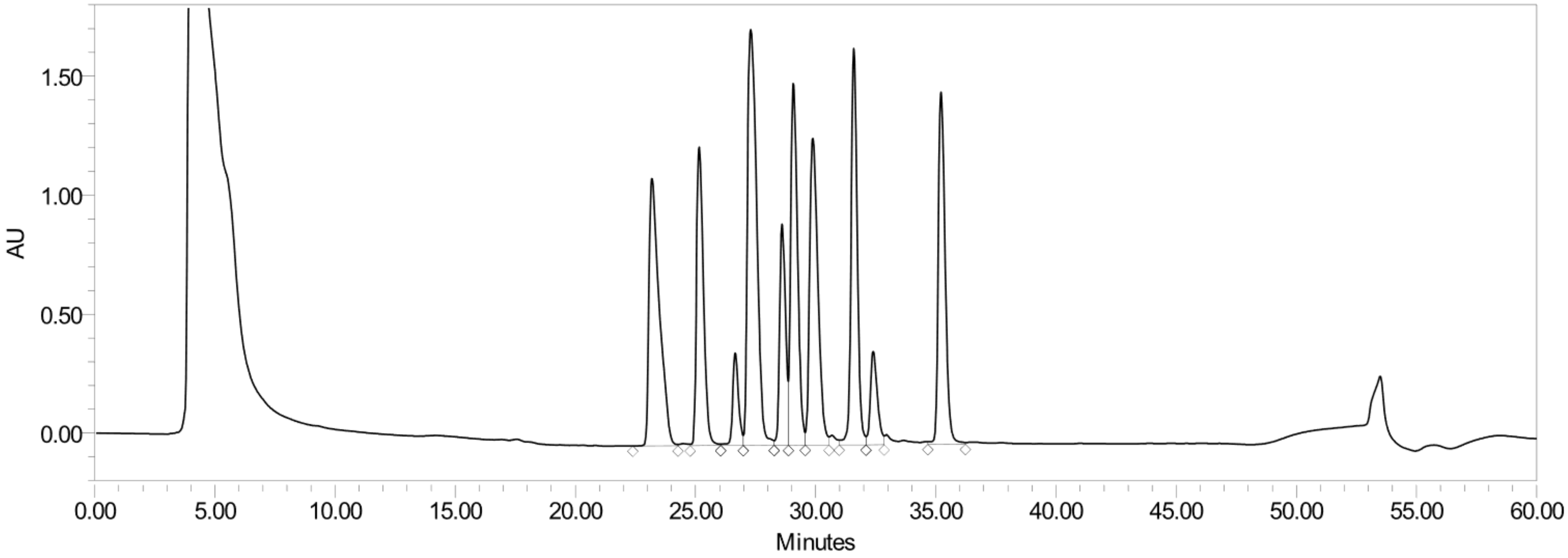
